# Quality Control Framework of TCM Preparations based on Multi-type Fingerprints using a Source Proportion Estimation Model

**DOI:** 10.1101/2022.04.27.489828

**Authors:** Yuguo Zha, Qi Yao, Dan Zhao, Xue Zhu, Hong Bai, Kang Ning

**Author notes:** Corresponding author., (Kang Ning), (Hong Bai).

## Abstract

Traditional Chinese Medicine (TCM) preparations have been widely used in clinical practice for the treatment of various diseases. The quality of TCM preparations is related to clinical efficacy and safety and is highly valued by researchers. The authenticity of TCM preparation can be guaranteed objectively by accurate quality control according to the composition. Here, we proposed a quality control framework of TCM preparations, which is based on multi-type fingerprints using the source proportion estimation model (SPEM). The high-performance liquid chromatography (HPLC) analysis and the high-throughput sequencing analysis are employed to acquire the chemical and taxonomic fingerprints of samples, respectively. The quality of TCM preparations among different manufacturers or batches is evaluated by using SPEM, which is an unsupervised method for source identification of TCM samples. Results showed the good performance of the quality control framework, for example, SPEM achieved a mean accuracy of 0.778 based on the ITS2 taxonomic fingerprint when differentiating manufacturer of BazhenYimu Wan pill. Applications of the quality control framework revealed the batch effect in TCM samples, and environmental factors, such as geography have a profound impact on the consistency of TCM preparations. In summary, this study is an exploration in the field of digital development of TCM preparations and provide a new insight to quantify the batch effect among different batches of TCM samples.

## Introduction

Traditional Chinese medicine (TCM) preparation has been widely used in clinical practice in China for tens of centuries^1-4^. However, the quality and safety of TCM preparations remain key concerns around the world, which hinder their broader application and popularity among international healthcare practitioners^5^. In recent years, due to the proven therapeutic effects of several authentic and precious TCM preparations, the adulteration, substitution and mislabeling of TCM become a global concern^6^. There are many reports on the traceability of Chinese medicinal materials based on DNA barcoding technology^7,8^. It is necessary to ensure the authenticity and reliability of TCM production by means of traceability. Quality control of TCM preparations, identifying from which manufacturer or batch the TCM (including the forms of pills, powders, capsules, tablets, etc.) is from, would be critical in TCM industry, for both producers and consumers alike. However, the quality control of TCM herbs is much more difficult than quality control of small molecules in western medicines. High quality TCM herbal preparations only comes from herbs with good quality, while the authenticity of TCM herbs is the first point of concern for quality.

TCM preparations are usually composed of several natural materials including plant, animal and mineral, based on which the therapeutic effects of TCM preparations are exerted. Therefore, the quality of TCM preparations is very important for clinical efficacy. For a long time, people relied on experience, from the appearance, smell and some simple physical and chemical phenomena of medicinal materials to judge their authenticity, but it is often very subjective and one-sided. With the development of modern molecular biology technology, the quality control methods of TCM have changed a lot. Quality control of TCM preparations recorded in Chinese Pharmacopoeia (Ch. P.) is mainly composed of the chemical ingredient (main chemical components) analysis and biological ingredient (taxonomy composition) analysis^9^. To date, studies focused on chemical ingredients of TCM preparations were abundant, while a few studies were reported for their biological ingredients. As an important part of TCM research, biological ingredients have drawn more and more attention around the world. Biological composition and chemical composition are inseparable parts for the quality control of TCM compound preparations. However, there is currently a lack of effective quality control framework of TCM preparations based on multi-type fingerprints^10^.

Here, we proposed a quality control framework of TCM preparations based on multi-type fingerprints using the source proportion estimation model (SPEM). The quality control framework is used to evaluate the quality of TCM preparations more accurately, comprehensively and systematically. The multi-type fingerprints include chemical fingerprint acquired by HPLC and taxonomic fingerprint acquired by high-throughput sequencing. HPLC is widely applied to characterize the chemical components in TCMs and is regarded as one of the most promising and reliable means for quality control of TCM preparations, and high-throughput sequencing is widely used to identify the taxonomy composition in biological samples. Multi-type fingerprints could be used to identify the sources and reflect changes in the intrinsic quality of TCMs. SPEM employs the source tracking method, FEAST^11^ to measure the similarities between TCMs samples. The combination of multi-type fingerprints with the source proportion estimation method can effectively discriminate TCMs from different geographical sources, parts, and cultivars and identify authenticity to prevent adulteration.

We used four types of TCM preparations as prototypes and performed experiments based on TCM samples from two manufacturers and three batches. Results showed the good performance of the quality control framework. For example, SPEM achieved a mean accuracy of 0.778 based on the ITS2 taxonomic fingerprint when identify which manufacturer the BazhenYimu Wan (BYW) pill sample is from. Applications of the quality control framework revealed the batch effect in TCM samples, and environmental factors, such as geography have a profound impact on the consistency of TCM preparations. In summary, this study is an exploration in the field of digital development of TCM preparations and provide a new insight to quantify the batch effect among different batches of TCM samples.

## Materials and Methods

### Sample preparations

#### Samples and Reagents

4 type of TCM preparations were purchased from 2 different Chinese manufacturers (namely A and B), and each with 3 batch numbers (I, II and III) (**Table 1**). Each batch was implemented with 3 parallel repeats, therefore there were 2*3*3 = 18 samples for each type of TCM preparation, and there were 4*18 = 72 samples in total.

**Table 1.**
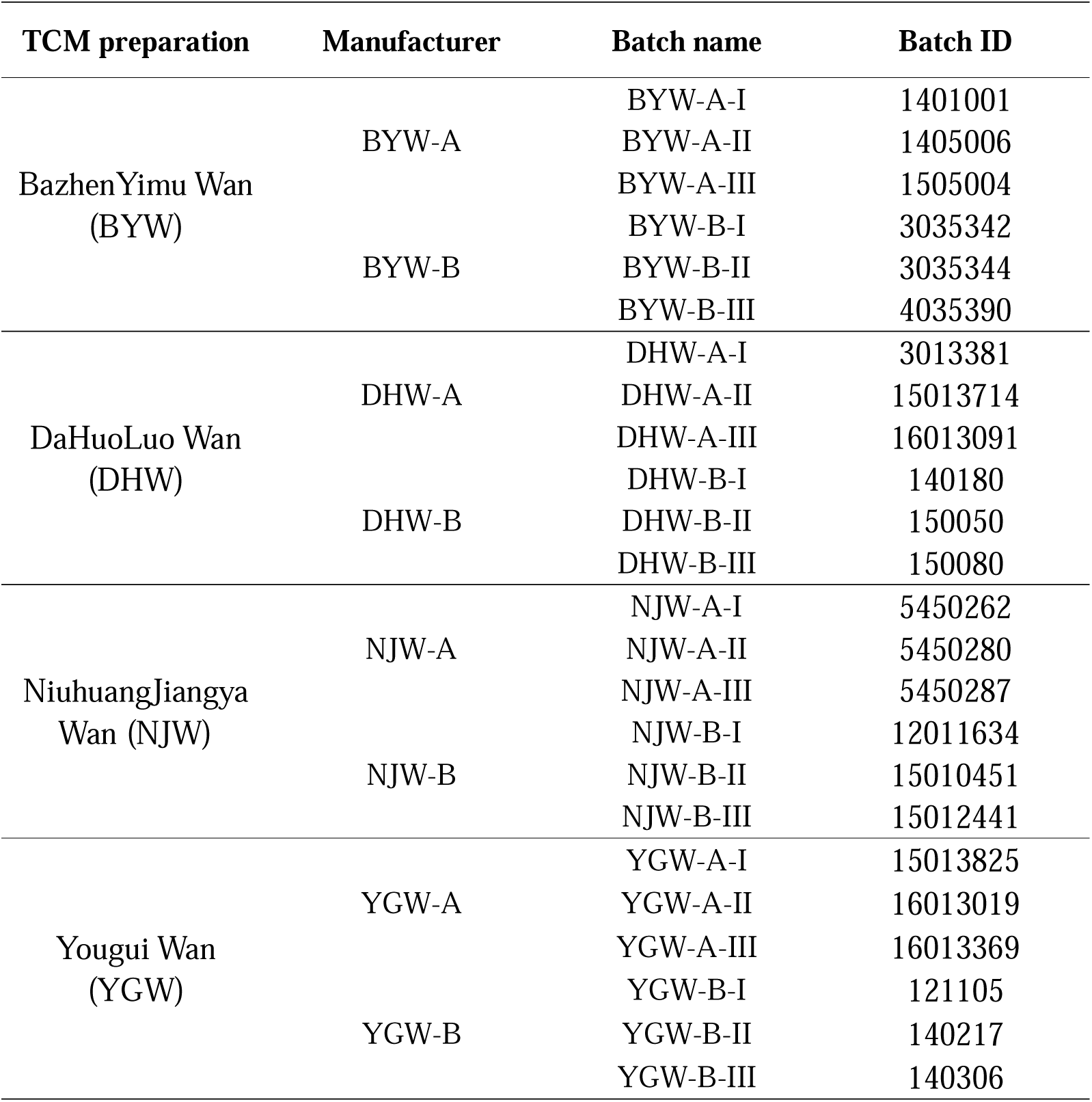
General information of samples used in this study.

| TCM preparation | Manufacturer | Batch name | Batch ID |
| --- | --- | --- | --- |
| BazhenYimu Wan<br>(BYW) | BYW-A | BYW-A-I | 1401001 |
|  |  | BYW-A-II | 1405006 |
|  |  | BYW-A-III | 1505004 |
|  | BYW-B | BYW-B-I | 3035342 |
|  |  | BYW-B-II | 3035344 |
|  |  | BYW-B-III | 4035390 |
| DaHuoLuo Wan<br>(DHW) | DHW-A | DHW-A-I | 3013381 |
|  |  | DHW-A-II | 15013714 |
|  |  | DHW-A-III | 16013091 |
|  | DHW-B | DHW-B-I | 140180 |
|  |  | DHW-B-II | 150050 |
|  |  | DHW-B-III | 150080 |
| Niu HuangJiangya<br>Wan (NJW) | NJW-A | NJW-A-I | 5450262 |
|  |  | NJW-A-II | 5450280 |
|  |  | NJW-A-III | 5450287 |
|  | NJW-B | NJW-B-I | 12011634 |
|  |  | NJW-B-II | 15010451 |
|  |  | NJW-B-III | 15012441 |
| Yougui Wan<br>(YGW) | YGW-A | YGW-A-I | 15013825 |
|  |  | YGW-A-II | 16013019 |
|  |  | YGW-A-III | 16013369 |
|  | YGW-B | YGW-B-I | 121105 |
|  |  | YGW-B-II | 140217 |
|  |  | YGW-B-III | 140306 |

These 4 types of TCM preparations include: BazhenYimu Wan (BYW), DaHuoLuo Wan (DHW), NiuhuangJiangya Wan (NJW) and Yougui Wan (YGW). In the recording of Chinese pharmacopoeia, their prescribed biological ingredients specified in Chinese pharmacopoeia were listed in (**Table S1** and **Table S2** in **Supplementary Materials**).

#### Sample preparation and analyses for biological ingredient

The steps of DNA extraction, amplification, sequencing and data analysis were described in our previous work^12,13^. Briefly, DNA was extracted by TCM-CTAB method, and these DNA extracts were amplified by touchdown PCR (by using ribosomal internal transcribed spacer 2, ITS2 and *trnL* as biomarker) before sent for Illumina MiSeq PE300 paired-end sequencing. After removing one *trnL*-marked BYW specimen that failed to be amplified and one ITS2-marked YGW sample that failed to be built the next-generation sequencing library preparation, the sequencing data of 142 samples was obtained and deposited and could be obtained from NCBI SRA database with accession number PRJNA562480.

Based on sequencing data, quality control, species identification and reads mapping of each species have been performed by following step: Reads from the same sample were assembled together by using ‘join_paired_end.py’ script in QIIME environment. Then the double-end barcodes (**Table S3** in **Supplementary Materials**) was extracted from all reads, and the ‘split_libraries_fastq.py’ was used to split the sample according their barcodes from the mixed sequencing data, and the Cutadapt command to remove the primers (**Table S4** in **Supplementary Materials**) from all samples. For every sample, the reads were then filtered by MOTHUR. We discarded <150 bp or >510 bp ITS2 reads, and <75 bp *trnL* reads. We also filtered the sequence whose average quality score was below 20 in each five bp-window rolling along with the whole reads. Then the sequences that contained ambiguous base call (N), homopolymers of more than eight bases or primers mismatched, uncorrectable barcodes, were also removed from all datasets. To match the most matched species for each sequence, we used the BLASTN (E-value=1E-10) to search in ITS2 and *trnL* database based on GenBank, respectively. Among all results, we first chose the prescribed herbal species with the highest score, else we selected the top-scored species. Then, we discarded the corresponding species of ITS2 and *trnL* sequences with relative abundance below 0.002 and 0.001, respectively.

### Chemical fingerprint processing

The chemical fingerprint of four kinds of TCM was established by high performance liquid chromatography (HPLC) method. The chemical fingerprint can reflect the characteristics of the TCM preparation to some extent, and can qualitatively compare the differences of chemical components in the TCM preparation, and then carry on the quality control of samples from different manufacturers and batches. Within a certain range, the content of a compound and its peak area are linear, which means that complex TCM preparations can be quantitatively identified by comparing the fingerprint feature. This also serves as the theoretical basis for the use of fingerprints in TCM preparations quality control^14,15^. Details about chemical fingerprints for these TCM preparations were provided in **Table S5** in **Supplementary Materials**.

### Taxonomic fingerprint processing

The identities and normalized relative abundances of species identified from TCM preparation samples represent the basic elements of the taxonomic fingerprint of samples. There are two types of taxonomic fingerprint (ITS2 and *trnL*) for each sample. For example, there were 41 ITS2 features and 72 *trnL* features for all of the 18 BYW samples. Details about taxonomic fingerprints for other preparations were provided in **Table S6** in **Supplementary Materials**.

### Source proportion estimation model and evaluation procedure

Source proportion estimation model (SPEM) employs the source tracking method (*i*.*e*., FEAST) to measure the similarities between TCMs samples. FEAST^11^ is an unsupervised method based on the Expected Maximization (EM) algorithm, which is successfully used in source tracking of microbial community samples. The great feature of FEAST is that it can tell in a sample with mixed species which proportions come from which different sub-environments. Here, we first consider two manufacturers as different sub-environments, and evaluate the source proportion of samples for each preparation and batch. Then, we consider three batches as different sub-environments, and evaluate the source proportion of samples for each preparation and manufacturer. To evaluate the ability of SPEM on identifying TCM samples from different manufacturers or batches, we used accuracy to represent such ability. For each TCM preparation (e.g., BYW) and each fingerprint (e.g., ITS2), TCM samples from the same batch (e.g., Batch-I) were selected for source identification via leave-one-out experiments. Specifically, there are 6 samples from the batch I of TCM preparation BYW, we used one sample (assume from manufacturer A) as unknown sink (i.e., query sample), and the remains (5 samples, 2 from manufacturer A and 3 from manufacturer B) as source samples. Then, SPEM was conduct to tell in the query sample of unknown sink which proportions come from which different sources (manufacturers). If the proportion of manufacturer A is the biggest proportion, then it is a correct case. For all the six samples, we performed six experiments, and count the accuracy as the number of correct cases over the number of total cases.

### SPD measurement

We defined the SPD score as source proportion divergence which is used for quantifying the batch effect among samples of TCM preparation. The SPD score is between 0 (no batch effect) and 1 (maximum batch effect). Specifically, for one preparation *p* and one manufacturer *m, SPD*_*p,m*_ score could be computed with the following formula:

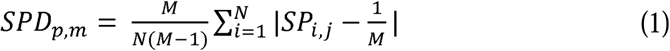

where *N* is the number of samples belong to preparation *p* and manufacturer *m, M* is the number of sub-environments (batches) involved, *SP*_*i,j*_ represent the source proportion of sample i from sub-environment *j*. For example, for preparation BYW and manufacturer A, there are 9 samples and 3 sub-environments (batches I, II and III).

For one preparation *p* and one batch *b, SPD*_*p,b*_ score could be computed with the following formula:

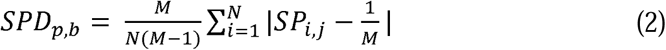

where *N* is the number of samples belong to preparation *p* and batch *b*, M is the number of sub-environments (manufacturers) involved, *SP*_*i,j*_, represent the source proportion of sample *i* from sub-environment *j*. For example, for preparation BYW and batch I, there are 6 samples and 2 sub-environments (manufacturer A and B).

## Results

### Quality control framework of TCM preparations

The newly proposed quality control framework of TCM preparations could be described as following workflow (**Figure 1**). First, chemical fingerprint and taxonomic fingerprint are obtained by high performance liquid chromatography (HPLC) analysis and high-throughput sequencing analysis, respectively. Second, SPEM employs the source tracking method, FEAST^11^, to measure the similarities between TCMs samples. Third, source proportion divergence (SPD, see **Materials and Methods**) is used to measure the batch effect in TCM samples.

**Figure 1.**
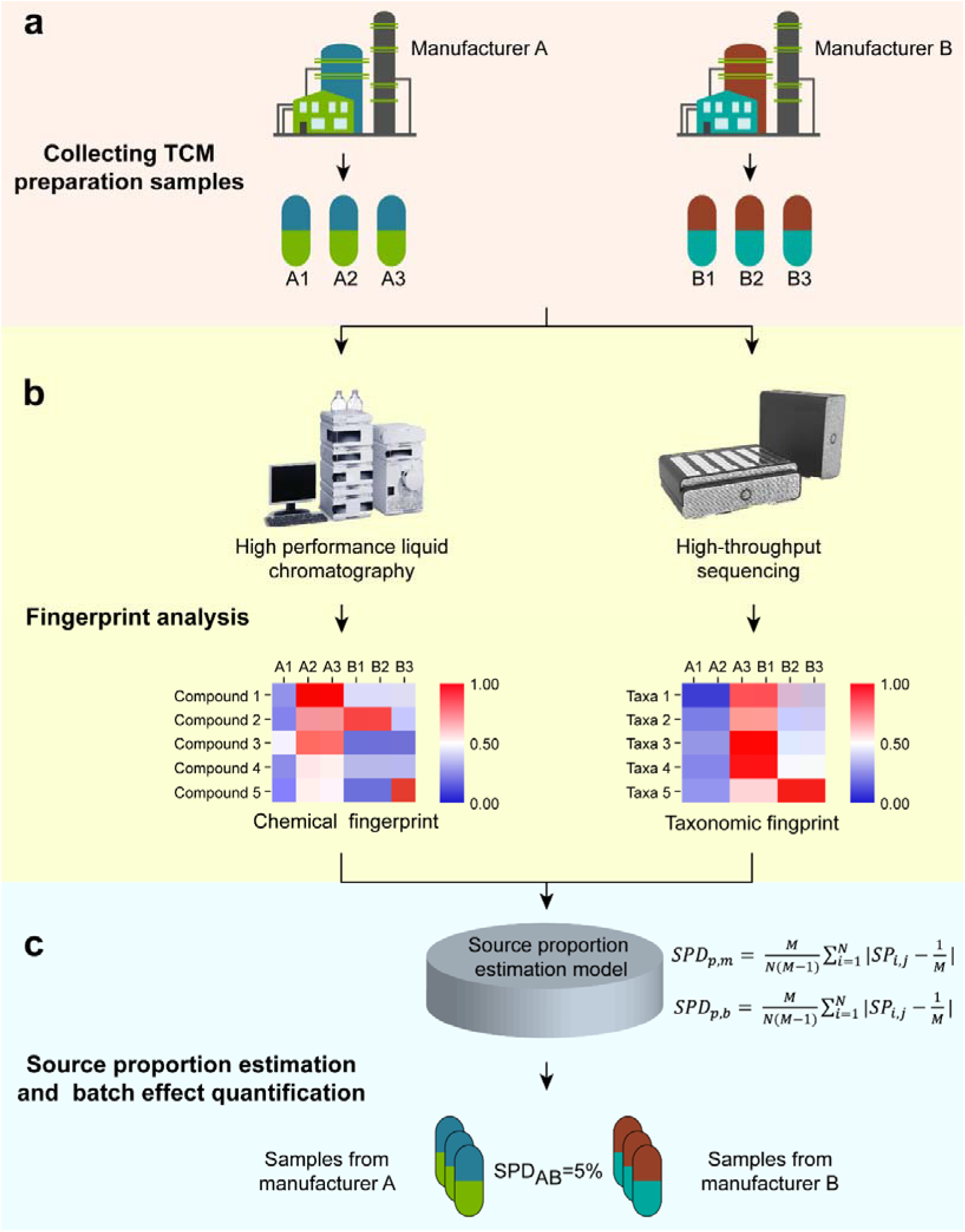
The newly proposed quality control framework of TCM preparations. **a, b**. TCM samples are collected and processed to produce the chemical and taxonomic fingerprints. **c**. Utilizing SPEM to measure the similarities between TCMs samples and measuring the batch effect in TCM samples with SPD. SPD, source proportion divergence.

### Similarity profiling of samples for different TCM preparations

Here, we took three batches of samples from two manufacturers with four types of TCM preparations as prototypes for testing (see **Materials and Methods**). The distance-based approach (i.e., Bray-Curtis) was first applied on all samples to provide an overview of the similarities among samples. We performed similarity profiling on the samples of four types of TCM preparations. Results of similarity profiling of samples for each preparation are showed in **Figure 2**.

**Figure 2.**
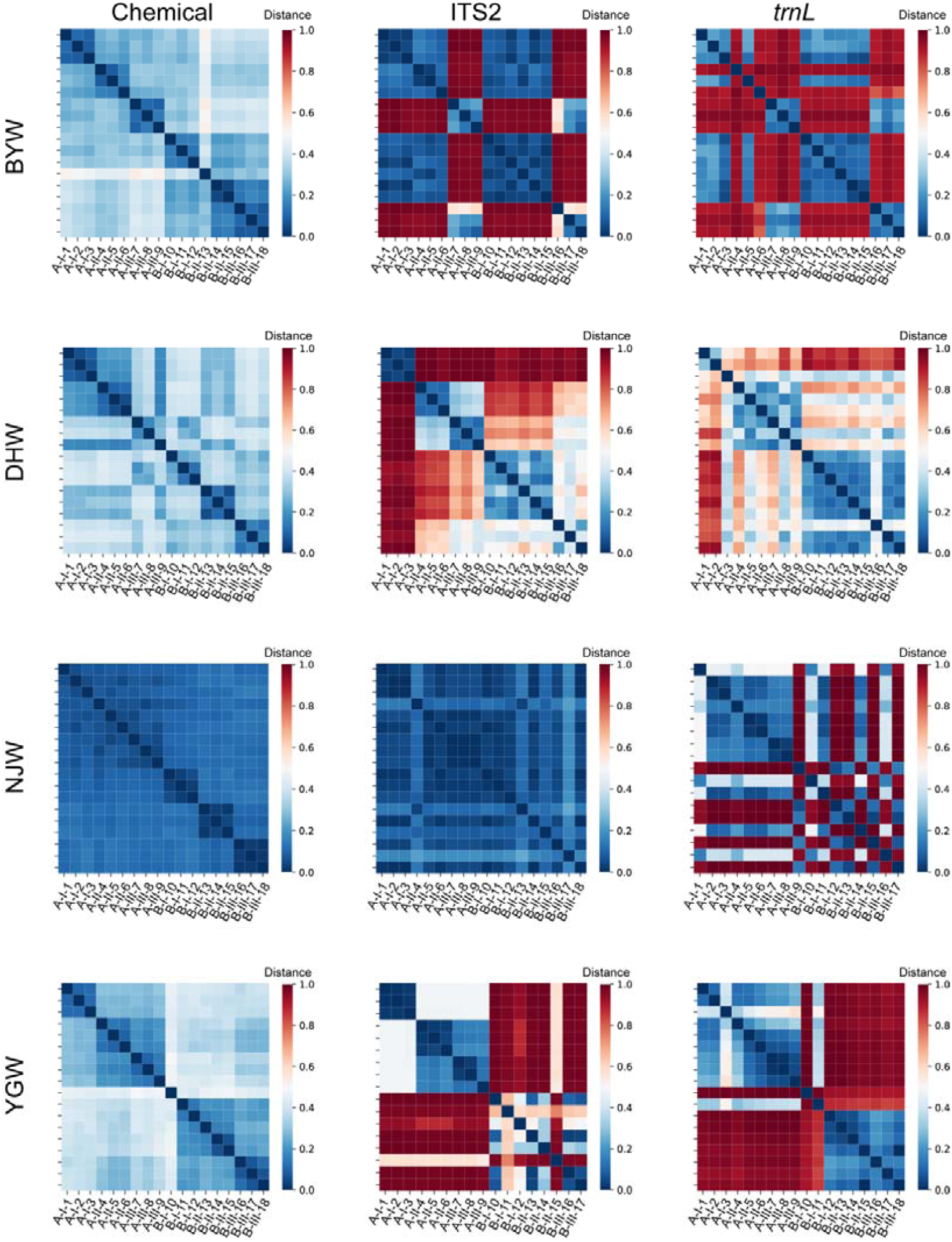
Similarity profiling of samples for each preparation based on chemical and taxonomic fingerprints. Color key indicates Bray-Curtis distance between TCM samples, which ranges from 0 to 1.

The Bray-Curtis distance among samples from different manufacturers but the same batch is relatively large, while Bray-Curtis distance among samples from different batches but the same manufacturer is relatively small. For example, the Bray-Curtis distance based on ITS2 taxonomic fingerprint among samples of YGW from two different manufacturers is relatively large (red color in batches I, II, and III), but the Bray-Curtis distance based on ITS2 taxonomic fingerprint among samples of YGW from three different batches is relatively small (blue color in manufacturers A and B). Here, we noticed that chemical fingerprint is more stable in measuring Bray-Curtis distances among samples than taxonomic fingerprint (both ITS2 and *trnL*). A possible explanation for such observation is that the stable content of chemical mineral components contained in qualified TCM preparations, regardless of the manufacturer and batch. However, it does not mean that the biological components in TCM preparations are as stable as the mineral components. Thus, it is necessary to assess the quality of TCM preparations based on both chemical and taxonomic fingerprints.

### Quality control of TCM preparations

Quality control of TCM preparations, identifying which manufacturer or batch the TCM sample (including the forms of pills, powders, capsules, tablets, etc.) is from would be critical in TCM industry. Here, SPEM was applied on identifying TCM samples from various manufacturers and batches. We conducted sample source search for four types of TCM preparations based on the chemical and taxonomic fingerprints, and evaluated the accuracy of the search.

Firstly, we evaluated the accuracy of identifying TCM samples from various manufacturers. In general, compared to accuracy based on taxonomic fingerprint, accuracy based on chemical fingerprint is higher regardless of TCM preparations and batches. For example, the accuracy based on chemical fingerprint for BYW is 1, but only 0.778 for ITS2 and 0.556 for *trnL*, respectively (**Table 2**). In terms of taxonomic fingerprint, the overall accuracy based on ITS2 is a little higher than the overall accuracy based on *trnL*, e.g., 0.778 vs. 0.556 for BYW and 0.722 vs.0.522 for NJW (**Table 2**).

**Table 2.**
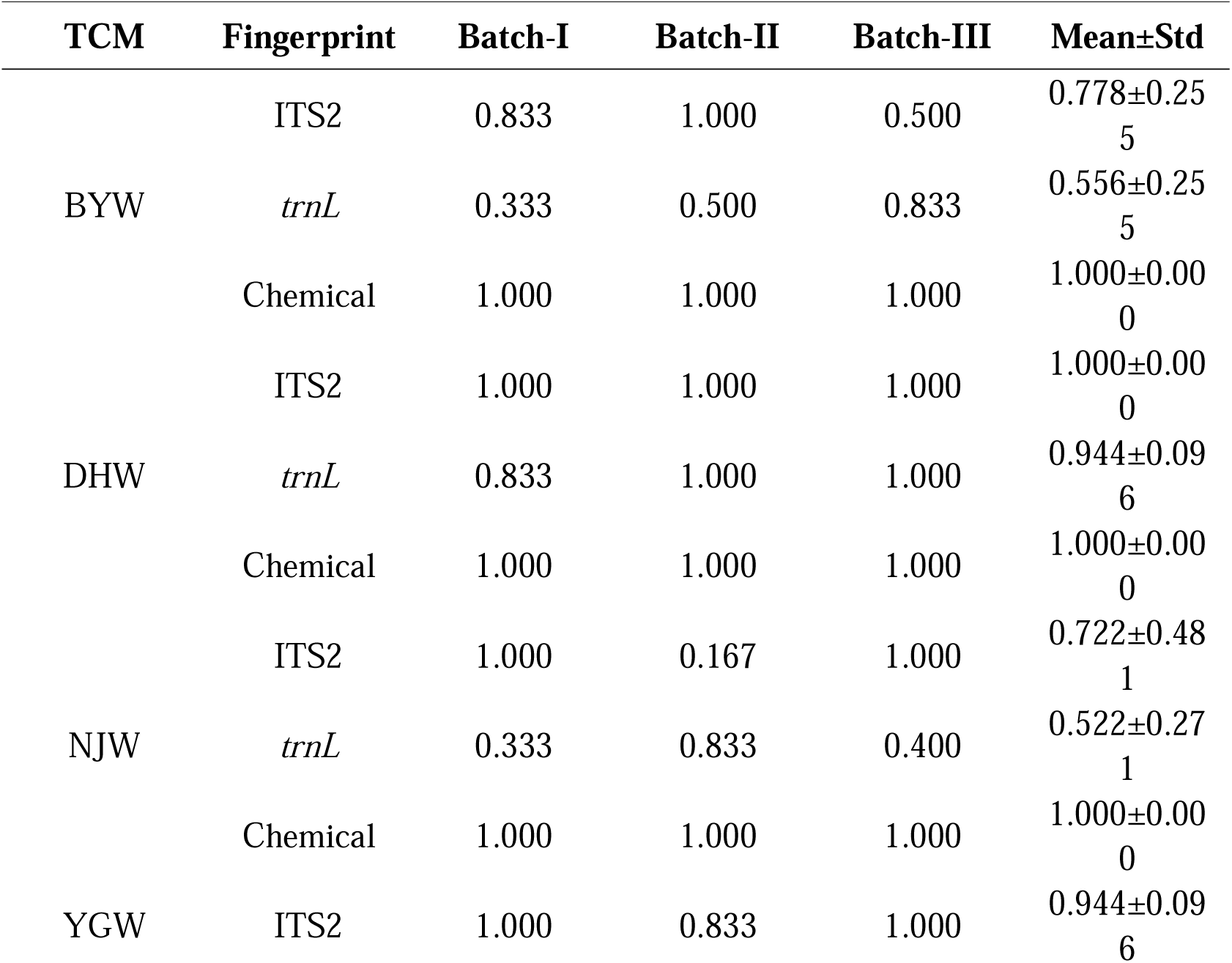
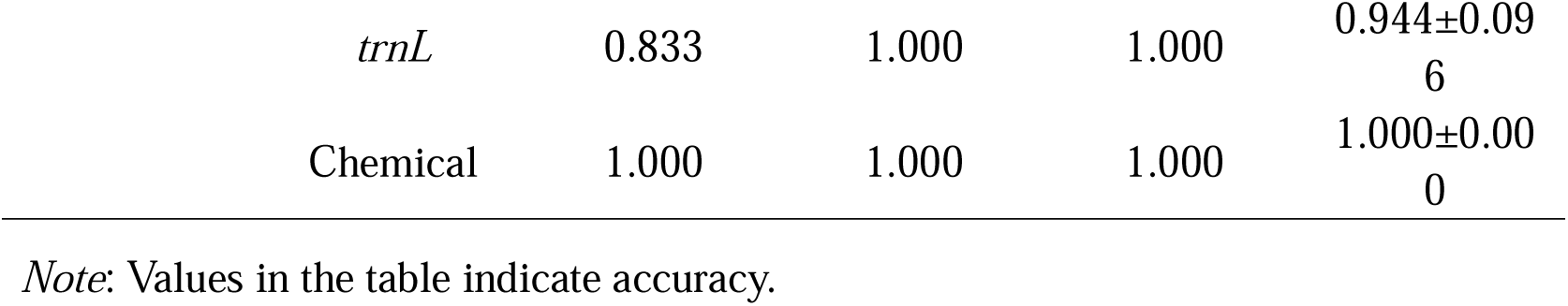
Accuracies of identifying TCM samples from various manufacturers using FEAST.

| TCM | Fingerprint | Batch-I | Batch-II | Batch-III | Mean±Std |
| --- | --- | --- | --- | --- | --- |
| BYW | ITS2 | 0.833 | 1.000 | 0.500 | 0.778±0.255 |
|  | *trnL* | 0.333 | 0.500 | 0.833 | 0.556±0.255 |
|  | Chemical | 1.000 | 1.000 | 1.000 | 1.000±0.000 |
| DHW | ITS2 | 1.000 | 1.000 | 1.000 | 1.000±0.000 |
|  | *trnL* | 0.833 | 1.000 | 1.000 | 0.944±0.096 |
|  | Chemical | 1.000 | 1.000 | 1.000 | 1.000±0.000 |
| NJW | ITS2 | 1.000 | 0.167 | 1.000 | 0.722±0.481 |
|  | *trnL* | 0.333 | 0.833 | 0.400 | 0.522±0.271 |
|  | Chemical | 1.000 | 1.000 | 1.000 | 1.000±0.000 |
| YGW | ITS2 | 1.000 | 0.833 | 1.000 | 0.944±0.096 |

|  |  |  |  |  |
| --- | --- | --- | --- | --- |
| *trnL* | 0.833 | 1.000 | 1.000 | 0.944±0.096 |
| Chemical | 1.000 | 1.000 | 1.000 | 1.000±0.000 |

*Note:* Values in the table indicate accuracy.

Secondly, we evaluated the accuracy of identifying TCM samples from various batches. In general, compared to accuracy based on taxonomic fingerprint, accuracy based on chemical fingerprint is higher regardless of TCM preparations and manufacturers. For example, the search accuracies based on chemical fingerprint for BYW is 0.944, but only 0.722 for both ITS2 and *trnL*, (**Table 3**). In terms of taxonomic fingerprint, accuracy based on ITS2 is a little higher than accuracy based on *trnL*, e.g., 0.778 vs. 0.444 for DHW and 0.667 vs. 0.157 for NJW (**Table 3**).

**Table 3.**
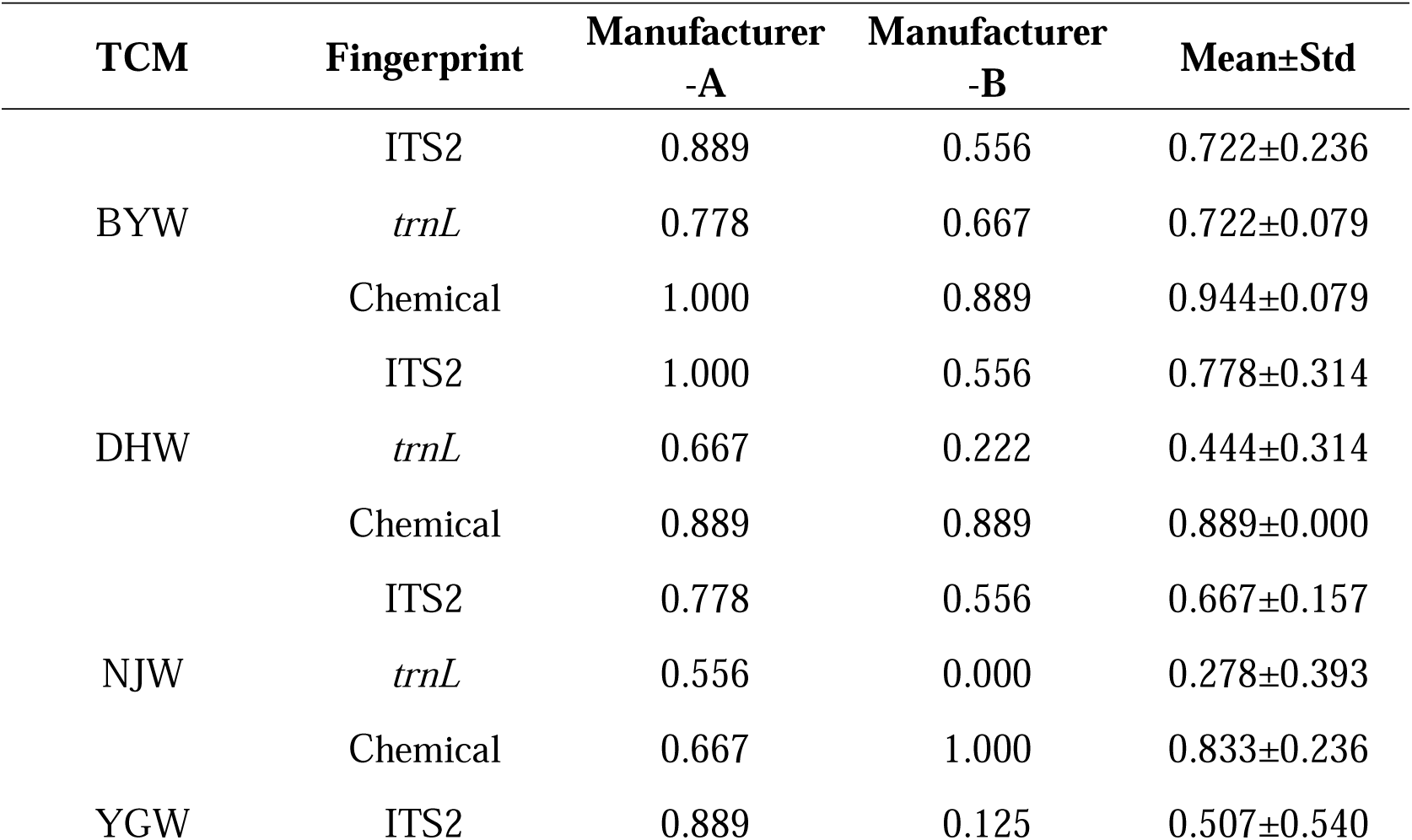
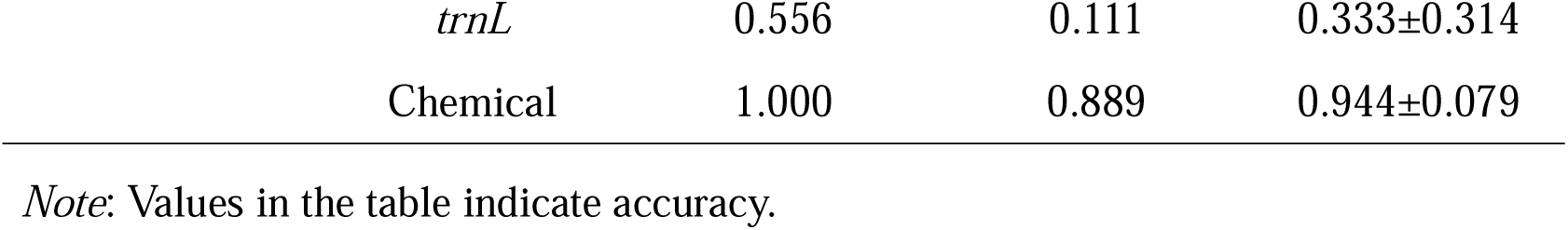
Accuracies of identifying TCM samples from various batches using FEAST.

| TCM | Fingerprint | Manufacturer<br>-A | Manufacturer<br>-B | Mean±Std |
| --- | --- | --- | --- | --- |
| BYW | ITS2 | 0.889 | 0.556 | 0.722±0.236 |
|  | *trnL* | 0.778 | 0.667 | 0.722±0.079 |
|  | Chemical | 1.000 | 0.889 | 0.944±0.079 |
| DHW | ITS2 | 1.000 | 0.556 | 0.778±0.314 |
|  | *trnL* | 0.667 | 0.222 | 0.444±0.314 |
|  | Chemical | 0.889 | 0.889 | 0.889±0.000 |
| NJW | ITS2 | 0.778 | 0.556 | 0.667±0.157 |
|  | *trnL* | 0.556 | 0.000 | 0.278±0.393 |
|  | Chemical | 0.667 | 1.000 | 0.833±0.236 |
| YGW | ITS2 | 0.889 | 0.125 | 0.507±0.540 |

|  |  |  |  |
| --- | --- | --- | --- |
| *trnL* | 0.556 | 0.111 | 0.333±0.314 |
| Chemical | 1.000 | 0.889 | 0.944±0.079 |

*Note:* Values in the table indicate accuracy.

### Batch effect evaluation based on source proportion divergence

We noticed that there are different degrees of batch effect in TCM samples for each TCM preparation, and batched effect existed in samples from two different manufacturers or samples from three different batches. We used source proportion divergence (SPD, see **Materials and Methods**) to quantify the batch effect that existed in samples from different manufacturers (**Table 4**) and batch effect that existed in samples from different batches (**Table 5**). The value of SPD is between 0 and 1, and the closer SPD is to 0, the smaller the batch effect is. On the contrary, the closer SPD is to 1, the larger the batch effect is.

**Table 4.** Source proportion divergence of TCM samples from different manufacturers.

| TCM | Fingerprint | Batch-I | Batch-II | Batch-III | Mean±Std |
| --- | --- | --- | --- | --- | --- |
| BYW | ITS2 | 0.164 | 0.220 | 0.190 | 0.192±0.028 |
|  | *trnL* | 0.189 | 0.191 | 0.200 | 0.193±0.006 |
|  | Chemical | 0.283 | 0.281 | 0.291 | 0.285±0.005 |
| DHW | ITS2 | 0.315 | 0.287 | 0.224 | 0.275±0.047 |
|  | *trnL* | 0.236 | 0.258 | 0.162 | 0.219±0.050 |
|  | Chemical | 0.271 | 0.290 | 0.219 | 0.260±0.037 |
| NJW | ITS2 | 0.222 | 0.199 | 0.287 | 0.236±0.045 |
|  | *trnL* | 0.192 | 0.303 | 0.288 | 0.261±0.060 |
|  | Chemical | 0.285 | 0.272 | 0.301 | 0.286±0.015 |
| YGW | ITS2 | 0.191 | 0.240 | 0.251 | 0.228±0.032 |
|  | *trnL* | 0.249 | 0.309 | 0.321 | 0.293±0.039 |
|  | Chemical | 0.232 | 0.283 | 0.284 | 0.267±0.030 |

*Note:* Values in the table indicate source proportion divergence.

**Table 5.**
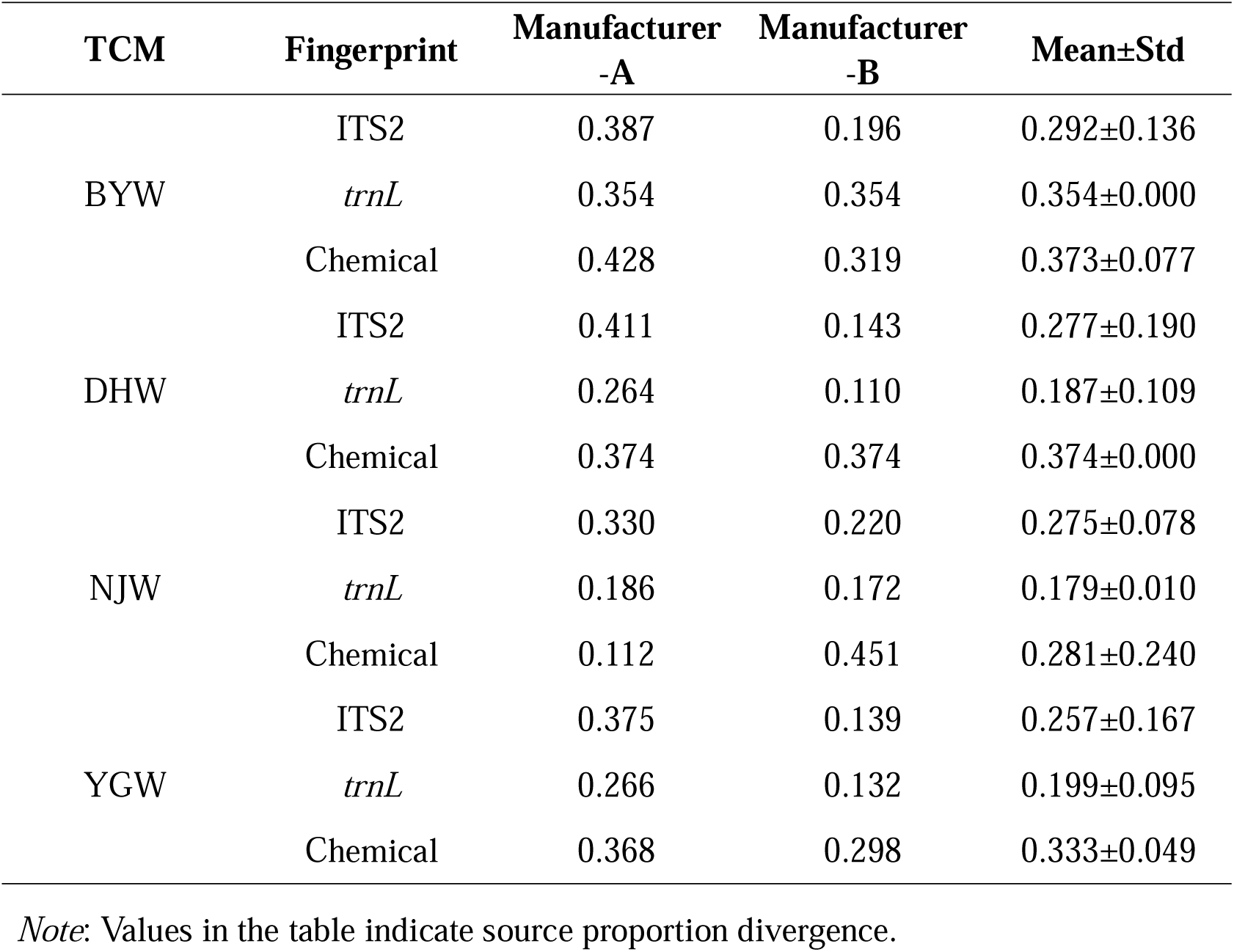
Source proportion divergence of samples from different batches.

| TCM | Fingerprint | Manufacturer<br>-A | Manufacturer<br>-B | Mean±Std |
| --- | --- | --- | --- | --- |
| BYW | ITS2 | 0.387 | 0.196 | 0.292±0.136 |
|  | *trnL* | 0.354 | 0.354 | 0.354±0.000 |
|  | Chemical | 0.428 | 0.319 | 0.373±0.077 |
| DHW | ITS2 | 0.411 | 0.143 | 0.277±0.190 |
|  | *trnL* | 0.264 | 0.110 | 0.187±0.109 |
|  | Chemical | 0.374 | 0.374 | 0.374±0.000 |
| NJW | ITS2 | 0.330 | 0.220 | 0.275±0.078 |
|  | *trnL* | 0.186 | 0.172 | 0.179±0.010 |
|  | Chemical | 0.112 | 0.451 | 0.281±0.240 |
| YGW | ITS2 | 0.375 | 0.139 | 0.257±0.167 |
|  | *trnL* | 0.266 | 0.132 | 0.199±0.095 |
|  | Chemical | 0.368 | 0.298 | 0.333±0.049 |

*Note:* Values in the table indicate source proportion divergence.

First, we investigated the batch effect among TCM samples from different manufacturers. Results showed an overall SPD values between 0.260 and 0.286 based on chemical fingerprint for those TCM preparations **(Table 4)**. However, the ITS2-based SPD values ranged from 0.192 to 0.275, and the *trnL*-based SPD values ranged from 0.193 to 0.293 (**Table 4**). Results suggested that there are different degrees of batch effect in samples for each type of TCM preparation. Moreover, results showed that the batch effect existing in two manufacturers based on chemical fingerprint was larger than the batch effect based on taxonomic fingerprint. For example, the SPD value based on chemical fingerprint of BYW is 0.285, which is significantly larger than the SPD value based on ITS2 or trn*L* fingerprint (i.e., 0.192 for ITS2, 0.193 for trn*L*).

Second, we investigated the batch effect among samples from different batches. Results showed the SPD values of manufacturer-B is always smaller than the SPD values of manufacturer-A (**Table 5**) and the batch effect existing in three batches based on chemical fingerprint was larger than the batch effect based on taxonomic fingerprint. For example, the SPD value based on chemical fingerprint of BYW is 0.373, which is significantly larger than the SPD value based on ITS2 or trn*L* fingerprint (i.e., 0,292 for ITS2, 0.354 for trn*L*).

In summary, there is a certain degree of batch effect existing in TCM samples from the two manufacturers and the three batches, underscoring the challenges of quality control of chemical and biological components in TCM preparation samples. SPEM revealed the relative stability of biological components in TCM preparation samples compared to chemical components. Notably, the batch effect among TCM preparation samples from two manufacturers suggested that the same TCM formula from different manufacturers might need further optimization^16^. While on the other hand, we can not exclude the possibility that different manufacturers have optimized TCM preparations for certain TCM formula so that they could be more suitable for medical treatment the local population^17,18^.

## Discussions

The quality of TCM preparations is related to clinical efficacy and safety, which is highly valued by people. TCM preparations consists of several herbs, which may contain hundreds or thousands of ingredients, which undoubtedly brings great difficulties to the quality control. Influenced by factors such as place of origin, growth time, harvest time, planting and processing technology, the quality of TCM preparations varies highly, leading to poor consistency of quality in different manufacturers and batches of preparations. Quality consistency has become a difficulty in the development of TCM industry, which profoundly affects the stable and controllable clinical efficacy of TCM and the repeatability and recognition of modern research results. However, rigorous quality control and assessment of components of TCM preparation are infrequently reported for TCM studies.

In this study, we proposed and evaluated a framework for an in-depth approach to a comprehensive quality control assessment of TCM preparation. The framework consists of three stages: (1) apply high-throughput sequencing analysis to obtain taxonomic fingerprint of TCM preparation samples, and apply HPLC analysis to obtain chemical fingerprint of TCM preparation samples; (2) utilizing SPEM to measure the similarities between TCMs samples; (3) measuring the batch effect in TCM samples with SPD. In this study, we took three batches of samples from two manufacturers with four types of TCM preparations as prototypes for testing and evaluation. Additionally, we also used this framework to quantitatively analyze the batch effect of TCM preparation samples. Results showed the good performance of the quality control framework and revealed the batch effect among TCM samples. Most importantly, the quality of TCM samples is stable on the premise of meeting the single dosage of each component, no matter the chemical or biological component.

In summary, the SPEM model and SPD measurement in combination are powerful for quality control of TCM preparations based on multi-type fingerprints. The integration of chemical fingerprint and taxonomic fingerprint revealed the chemical and biological characteristics of different TCM preparations, and SPEM was proved to be successful for quantifying these differences. While SPD could further quantify the batch effects of samples from different manufacturers and batches. Applications of the quality control framework on four types of TCM preparations showed the ability of the framework on both source identification and batch effect quantification in TCM samples. This would not only be of values for large-scale TCM preparation screening, but also could be used in clinics for quick and reliable source tacking.

## Conclusion

Taken together, our study utilized multi-type fingerprints and SPEM model, which could help for explaining the relationship between these ingredient variations and the authenticity of TCM preparations. This study is an explorative study in the field of digital development of TCM preparations, illustrates the quantification platform for TCM preparation quality control, offers a new insight to quantify the batch effect among different batches of TCM samples, and provides future perspectives of using both chemical and taxonomic fingerprints combined with unsupervised/supervised methods towards accurate and fast quality control of TCM preparations.

## Declarations

### Ethics approval and consent to participate

Not applicable

### Consent for publication

Not applicable

### Availability of data and materials

The datasets generated and/or analyzed during the current study are available in the NCBI Sequence Read Archive (SRA) repository with accession number PRJNA562480.

### Authors’ contributions

H.B. and K.N. conceived of and proposed the idea, and designed the study. Y.Z., Q.Y. and X.Z. performed the experiments. H.B., Y.Z, D.Z., X.Z. and K.N. analyzed the data. H.B., Y.Z., D.Z., X.Z. and K.N. contributed to editing and proof-reading the manuscript. All authors read and approved the final manuscript.

## Acknowledgments

We thank GenWiz Inc. for conducting the high-throughput sequencing of the samples.

